# Population-scale proteome variation in human induced pluripotent stem cells

**DOI:** 10.1101/439216

**Authors:** Bogdan A Mirauta, Daniel D Seaton, Dalila Bensaddek, Alejandro Brenes, Marc J Bonder, Helena Kilpinen, HipSci Consortium, Oliver Stegle, Angus I Lamond

## Abstract

Realising the potential of human induced pluripotent stem cell (iPSC) technology for drug discovery, disease modelling and cell therapy requires an understanding of variability across iPSC lines. While previous studies have characterized iPS cell lines genetically and transcriptionally, little is known about the variability of the iPSC proteome. Here, we present the first comprehensive proteomic iPSC dataset, analysing 202 iPSC lines derived from 151 donors. We characterise the major genetic determinants affecting proteome and transcriptome variation across iPSC lines and identify key regulatory mechanisms affecting variation in protein abundance. Our data identified >700 human iPSC protein quantitative trait loci (pQTLs). We mapped *trans* regulatory effects, identifying an important role for protein-protein interactions. We discovered that pQTLs show increased enrichment in disease-linked GWAS variants, compared with RNA-based eQTLs.

## Introduction

Induced pluripotent stem cells (iPSC) hold enormous promise for advancing basic research and biomedicine. By enabling the *in vitro* reconstitution of development and cell differentiation, iPS cells allow the investigation of mechanisms underlying development and the aetiology of many forms of genetic disease. To realize this potential, it is essential to characterize how genetic and non-genetic effects in human iPSCs influence molecular and cellular phenotypes.

Recently, the establishment of population reference panels of normal human iPSC lines^1–3^ have provided valuable resources for functional experiments in different genetic backgrounds. Additionally, these data have yielded detailed characterizations of the iPS transcriptome, identifying thousands of *cis* expression Quantitative Trait Loci (eQTL)^1,4,5^, including at disease-relevant loci. While these RNA-based analyses are informative for studying mechanisms affecting gene regulation at the transcriptional level, most cellular phenotypes involve mechanisms acting downstream, at the protein level. Evidence in other contexts, including in lymphoblast cell lines and in cancer, point to substantial differences in the genetic regulation of protein and RNA traits, identifying protein QTL^6–9^ and assessing the extent of buffering of genetic effects between layers^10,11^. However, existing protein datasets have been limited by scale (i.e. number of samples) or resolution (i.e. number of proteins, availability of RNA data). Importantly, no population-scale proteome datasets have been generated from human pluripotent cells.

Here, we report on the first comprehensive, population-scale, combined proteomics and gene expression analysis in human iPSC lines. Our data comprise matched quantitative proteomic (TMT Mass Spectrometry) and transcriptomic (RNA-seq) profiles of 202 iPSC lines, derived from 151 donors that are part of the HipSci project^1^. We identify both genetic and non-genetic effects causing variability in protein expression between individuals. Our data provide the first high-resolution map of protein quantitative trait loci (pQTLs) in human iPSCs, which we characterise in relation to regulatory variants that affect the iPSC transcriptome. This reveals important roles for protein-protein interactions in propagating and buffering genetic effects on the human proteome. Additionally, we identify pQTLs linked to GWAS loci, underlining the importance of direct protein measurements for the characterisation of disease mechanisms.

## Results

### A population reference proteome for human iPSCs

We selected 217 iPSC lines from the HipSci project^1^, which were derived from 163 different donors, for protein analysis. Quantitative mass spectrometry was carried out in batches of 10, using tandem mass tagging (TMT^12^), including one common reference iPSC line that was included in each batch (**Methods**). After quality control (**Supp. Fig. 1; Methods**), we selected 202 lines (from 151 donors) for which genotype, RNA-seq and proteome information is available, for further analysis (**Fig. 1A**; **Supp. Table 1**).

**Figure 1.**
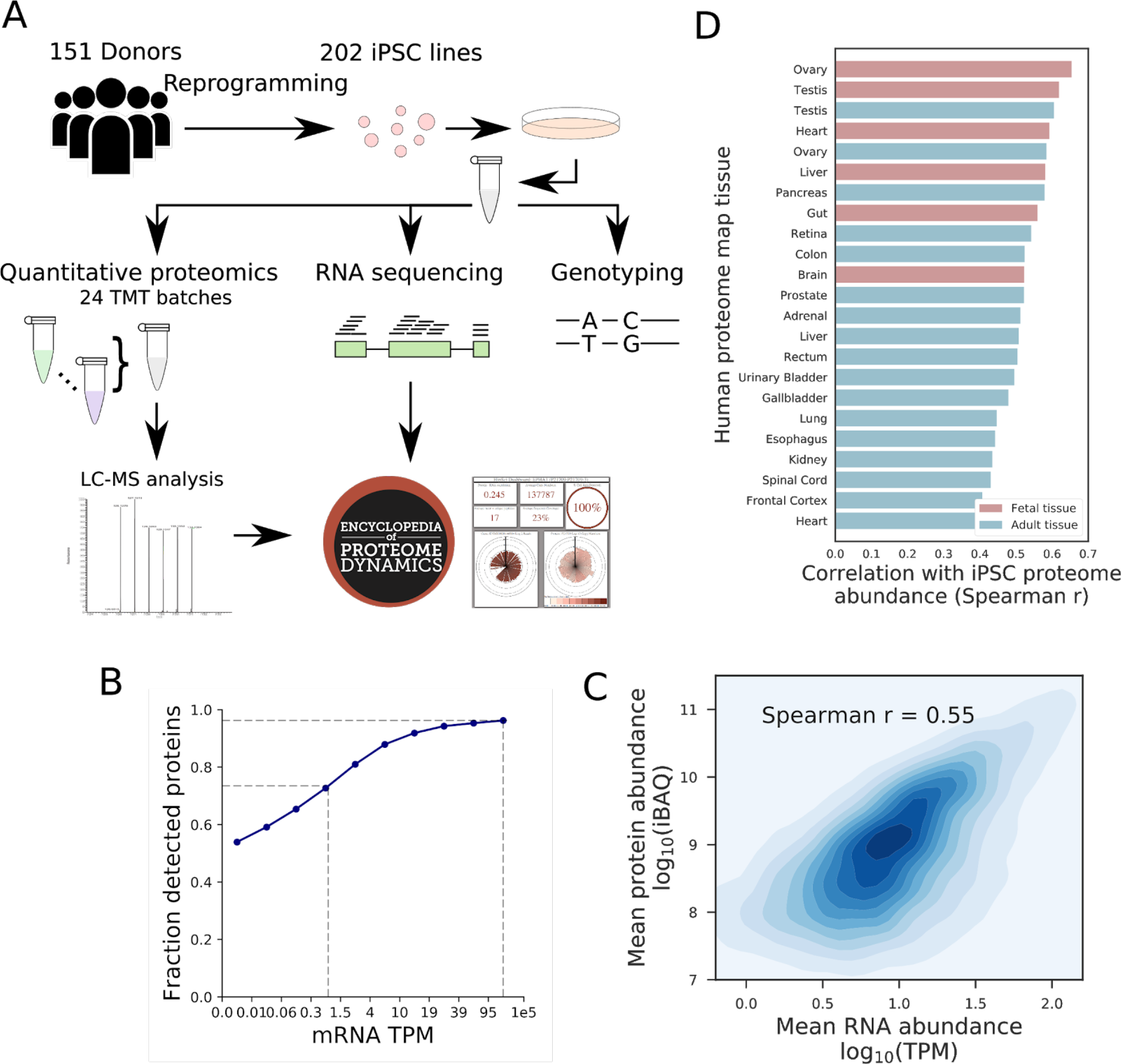
Molecular profiling of iPSC lines. **(A)** Experimental design, displaying assays considered in this study. Genotype, RNA-seq and quantitative proteomics data were generated from the same cell lines. **(B)** Aggregate proteome coverage, displaying the fraction of genes with detected protein peptides as a function of RNA abundance (mRNA transcripts per million reads). **(C)** Genome-wide correlation between the aggregate RNA and protein abundance for 10,672 protein-coding genes (showing average expression across 202 lines). All proteomics data can be interactively explored in the Encyclopedia of Proteome Dynamics (http://www.peptracker.com/epd). **(D)** Similarity between the iPSC proteome and somatic tissues. Shown are Spearman correlation coefficients between the average iPSC proteome and 23 tissues from the Human Proteome Map, including Adult (Red) and Fetal (Blue) tissues (**Methods**).

In aggregate, our proteomics data identified >250,000 distinct (unmodified) peptide sequences, corresponding to 16,218 protein groups (hereon denoted proteins) with a median sequence coverage of 46% (**Supp. Table 2**), and that map to 10,394 protein coding genes. Of these, 11,542 protein groups corresponding to 9,993 genes were detected in more than 30 lines and were considered for downstream analysis (**Supp. Fig. 2**). RNA-seq data from the same iPSC lines identified 12,363 expressed protein-coding genes (TPM>1), ~75% of which had evidence for expression at the protein level (**Fig. 1B**; **Supp. Fig. 3**). The average abundance for cognate protein and RNA expression in iPSCs was positively correlated across genes (**Fig. 1C**), consistent with observations in other cell types and organisms ^13,14^.

Our data provide the most comprehensive analysis of the human iPSC proteome reported to date, and one of the most comprehensive proteomic datasets reported for any human primary or derived cell type (**Supp. Table 3**). Comparison of iPSC lines derived from both healthy and disease bearing donors (**Supp. Table 4**), indicates no substantial global disease-linked differences, at either proteome or transcriptome levels (**Supp. Fig. 4**). Notably, when we compared the iPSC proteome with the Human Proteome Map ^15^, foetal and reproductive organs were identified as the tissues with the most similar protein expression patterns to iPS cells (**Fig. 1D**). This is consistent with the expression of pluripotency markers in foetal testis and ovaries ^16,17^.

### RNA and proteome variability

Across iPSC lines, the majority of genes showed low RNA and protein coefficients of variation (**Fig. 2A**), with only weak to moderate global correlation across the lines (**Fig. 2B**). Notably, many highly variable RNAs showed low covariation with protein (985 RNA-protein pairs with r < 0.2), indicating that the variation in protein abundance between iPSC lines is not explained solely by variation in RNA expression levels.

**Figure 2.**
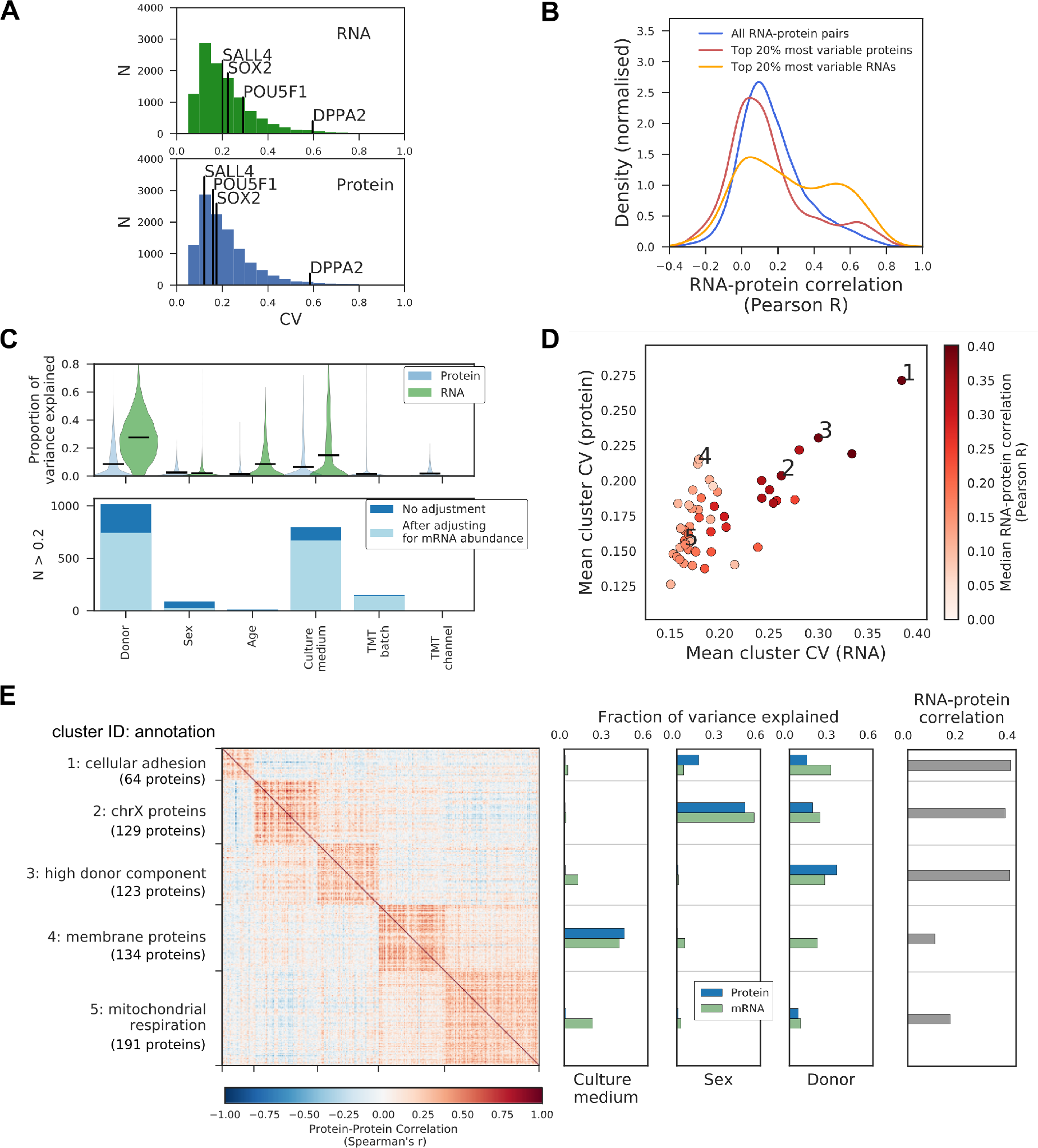
Genetic and non-genetic sources of iPS proteome variation. **(A)** Distribution of RNA and protein coefficients of variation for individual protein coding genes across lines. **(B)** Distribution of RNA-protein correlation coefficients for individual genes across lines (pearson r). Shown are densities for all genes or when selecting the top 20% variable RNA or protein. **(C)** Quantified variance components of individual RNA and protein, considering different technical and biological factors. Shown is the distribution of variance contributions of different factors (upper panel), and numbers of proteins with greater than 20% explained variance for each factor (lower panel). Also shown are the number of proteins that retain greater than 20% contribution for each factor when accounting for RNA variation (light blue; see **Methods**). **(D)** Median variability and RNA-protein correlations (Spearman r) across 51 protein co-expression modules. Specific modules of interest are labelled (1-5). **(E)** Left: Coexpression heatmap for proteins in modules labelled in **D**, displaying pairwise correlation coefficients between proteins (**Supp. Table 17**). Right: Variants components for the median protein and RNA levels of each module, as well as pairwise correlation (Pearson r).

Next, we assessed a range of factors, including the cell line donor, age, sex, as well as culture medium and other technical factors, for their potential contribution to the variation in protein expression between iPSC lines (**Fig. 2C**; **Methods**). The largest effects on protein variation were associated with donor effects and culture medium (**Fig. 2C**). Even after accounting for protein variability that can be explained by transcriptional mechanisms, i.e. where there was parallel variation in RNA expression (**Supp. Fig. 7**), substantial effects on protein expression levels were still observed for both donor and culture medium (**Fig. 2C**; **Methods**). This indicates that (i) differences between individual donors play an important role in causing the observed variation in proteome expression between the iPSC lines and (ii) post-transcriptional mechanisms also contribute significantly to these donor effects.

We note that some of the genes showing the strongest effect of donor variation on protein expression levels encoded the same proteins that were previously identified as being differentially expressed between reprogrammed iPS cells and embryonic stem cells (ESCs) ^18,19^. These earlier studies had suggested that reprogrammed iPS cells may have important differences in protein expression, when compared with the physiological stem cells present in embryos. However, these previous comparisons of iPSC and ESC cells did not control for genetic differences between donors. Our data show that these previously reported differences between iPSC and ESC cells may be explained by underlying effects of genetic variation between donors, rather than intrinsic differences between the iPSC and ESC cell types (**Supp. Fig. 6**). This supports the view that it is possible to reprogram iPS cells to a state showing near identical protein expression patterns to ESC cells.

### Coordinated expression changes of biological processes

Next, we explored protein co-expression clusters (**Methods**), which identified 51 modules of proteins that showed patterns of co-expression, 34 of which were enriched for at least 10 GO terms (FDR<10%; Fisher’s exact test; **Supp. Table 5**). Among the most prevalent processes identified were ‘cellular developmental process’ (3 modules), ‘cell adhesion’ (3 modules), and ‘respiratory electron transport chain’ (3 modules). For each module we evaluated: (i) the coefficients of protein abundance variation (CV), (ii) the fraction of variance explained by biological and technical factors (**Methods**), and (iii) the RNA-protein correlation. While modules with high protein variability also tended to show high RNA variability, (**Fig. 2D,E**; **Supp. Fig. 8**), we also identified clusters showing high variability at the protein level, but not at the RNA level (**Fig. 2D**).

There were differences between modules with high RNA and protein variability (e.g. clusters 1, 2, 3), both in the specific enriched GO terms and in their variance components (**Fig. 2D,E**). For example, Cluster 2 was enriched for proteins encoded on the X chromosome and variation was associated with the sex of the donor, at both the RNA and protein levels. In contrast, Cluster 4 showed high variability in protein abundance, but low RNA variability (**Fig. 2E**). The 134 proteins in Cluster 4 were enriched for integral membrane proteins and their variation was linked to the culture medium variance component (**Fig. 2E**), which was not explained by biases in the quantification of peptides from membrane proteins (**Supp. Fig. 5**). This indicates that differences in the cellular environment can affect the abundance of Cluster 4 proteins, and is not driven by changes in transcriptional regulation.

In summary, analysis across the 202 iPSC lines shows significant donor-to-donor variation in both the proteome and transcriptome. Interestingly, donor variation was apparent both at the level of individual proteins and in the coordinated regulation of whole pathways.

### Mapping *cis* genetic effects on protein abundance

Next, we mapped *cis* quantitative trait loci at both the RNA and protein levels (on autosomes; MAF>5%; within +/− 250 kb around the gene; using a linear mixed model; **Methods**). The number of pQTLs identified was greatly increased by adapting PEER adjustment to account for non-genetic sources of variation previously developed for mapping of RNA ^20^ to protein traits (**Methods**; **Supp. Fig. 10**). Proteomic QTL analysis identified 712 genes with a pQTL (FDR<10%; 10,675 proteins tested corresponding to 9,564 genes), compared to 5,744 genes with an eQTL when considering RNA levels (14,148 protein-coding and non-coding genes tested; 3,641 genes tested at both protein and RNA level; **Fig. 3A**; **Supp. Table 7,8,9**).

**Figure 3.**
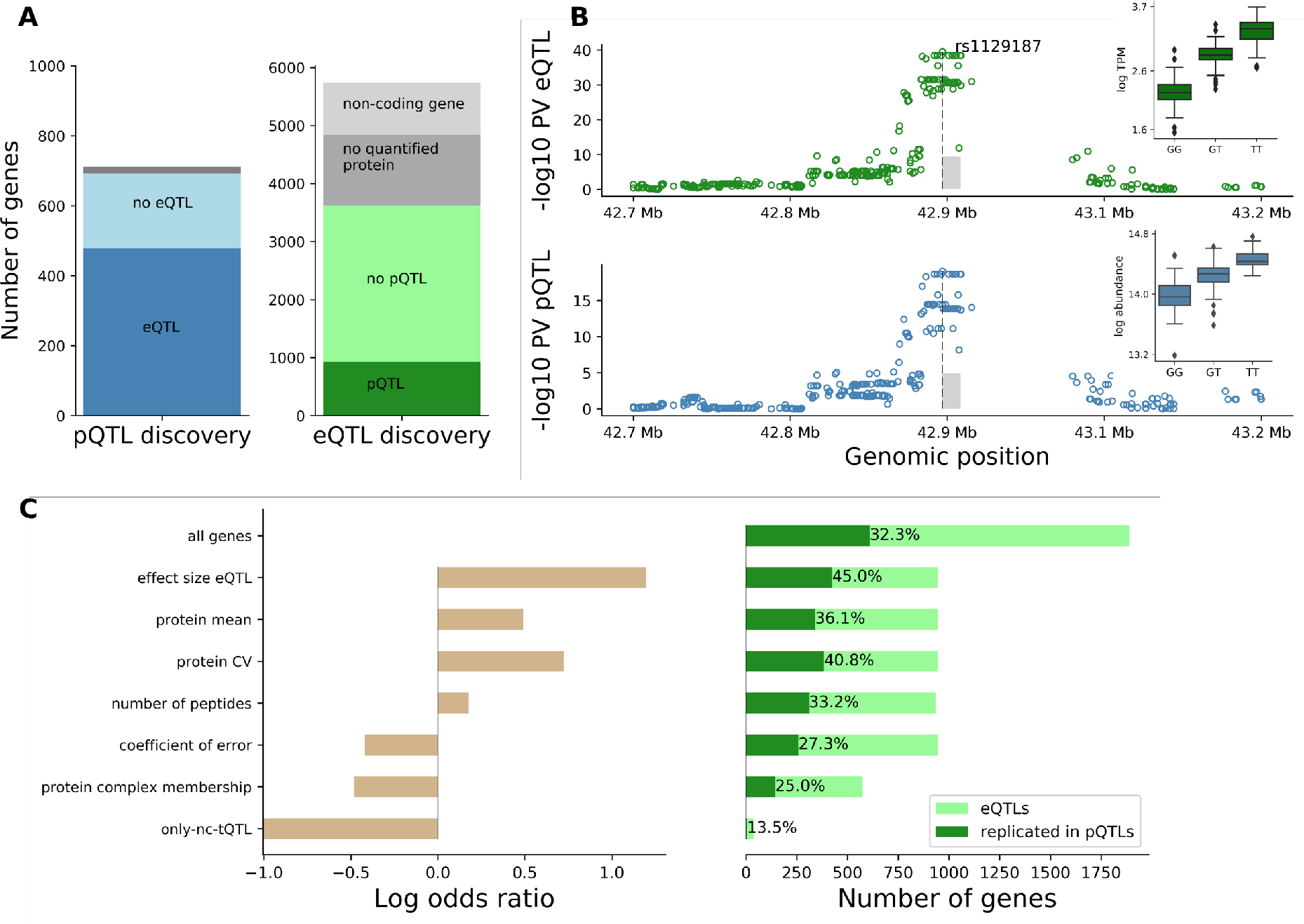
human iPSC *cis* protein and RNA QTLs. **(A)** Number of genes with a protein (blue) or RNA (green) QTL (FDR<10%) and replicated effects across molecular layers. Left: Number of pQTL genes, either with (dark blue) or without (light blue) replicated RNA effect. Right: Number of eQTL genes, either with (dark green) or without (light green) replicated protein effect. Replication defined at nominal P<0.01 with consistent effect direction. Grey fractions correspond to genes that could not be assessed at the other molecular layer (dark grey: not expressed, light grey: non-coding eQTL genes). (**B**) Manhattan plots for *cis* RNA (top) and protein (bottom) QTL mapping for *PEX6*. Boxplots show RNA and protein expression for different alleles at the eQTL lead variant rs1129187. **(C)** Logistic regression model trained on the replication status of eQTL at the protein level (defined as in **A**) considering technical and biological covariates (trained on 1,887 genes detected at protein and RNA level in all 202 lines; **Methods**). Left: Log odds ratio of individual covariates considered in the model. Right: Fraction of eQTLs with replicated protein effects, considering different gene strata. All genes correspond to no stratification. Considered covariates are: are eQTL effect size, average protein abundance, protein coefficient of variation across lines, number of identified protein peptides, protein technical coefficient of variation, membership in protein complexes, and whether the eQTL variant is associated with changes in expression of at least one coding transcript isoforms (only-nc-tQTLs). Percentages denote the replication rate.

To investigate which DNA sequence variants affected both protein and RNA expression levels, we assessed the ‘replication’ of pQTLs at the RNA level and vice versa (nominal significance at P<0.01 and same direction of effects; **Methods**). This revealed 478 pQTLs (69%) that were also detected at the RNA level. Conversely, analysis of 3,641 protein-coding eQTL genes with protein expression identified 897 eQTLs (25%) that were also detected at the protein level. Globally, eQTL and pQTL effect sizes were moderately correlated (**Supp. Fig. 11**). An example of an eQTL with a corresponding effect at the protein level is the lead eQTL variant rs1129187 for the *PEX6* gene (**Fig. 3B**), a known risk variant for Alzheimer’s disease in APOE e4+ carriers ^21^.

Next, we used multivariate logistic regression to systematically characterize the technical and biological determinants affecting whether eQTLs result in detectable protein changes **(Fig. 3C).** This identified the eQTL effect size as the most relevant positive factor, followed by the protein coefficient of variation and the average protein abundance (**Fig. 3C**; **Supp. Table 12, 13**). eQTLs for genes that are the subunits of protein complexes were less frequently detectable at the protein level. Notable examples include subunits of the mitochondrial ribosome and of the spliceosome, for which the eQTLs, while having highly significant effects at the RNA level, were buffered at the protein level (**Supp. Table 14**). This indicates that *cis* regulatory genetic effects on protein abundance in iPSCs can be tempered by post-transcriptional mechanisms dependent on protein-protein interactions. For comparison, we also considered technical sources of variation at the protein level (coefficient of error), which were markedly less relevant than biological factors. Therefore, we propose that the observed buffering of eQTLs at the protein level primarily arises from a combination of biological factors, rather than technical limitations in protein quantification.

### Isoforms affect eQTLs acting at the protein level

Next, we investigated the utility of RNA and protein quantification with isoform resolution to explain which eQTLs manifest in detectable protein effects. For this analysis we considered 54,965 transcript isoforms (quantified using Salmon^22^) and 126,758 peptides for QTL mapping, which identified 5,734 genes with a transcript QTL and 740 genes with a peptide QTL (**Supp. Fig. 13, Methods, Supp. Table 4,10,11**). Overlaying the iPSC transcript QTLs with gene-level eQTLs identified 84 eQTLs that were exclusively associated with abundance changes of a non-protein coding transcript isoform (nominal P<0.01). QTL analysis with transcript isoform resolution thus explains why some of the eQTLs identified by conventional RNA analysis cannot give rise to protein QTLs (**Fig. 3C**). For example, rs2709373, an eQTL variant for *METTL21A*, was associated specifically with the abundance of the non-coding transcript isoform ENST00000477919, without any detectable effect on the abundance of any protein-coding transcript isoforms and thus did not alter protein expression levels from this locus (**Fig. 4A**).

**Figure 4.**
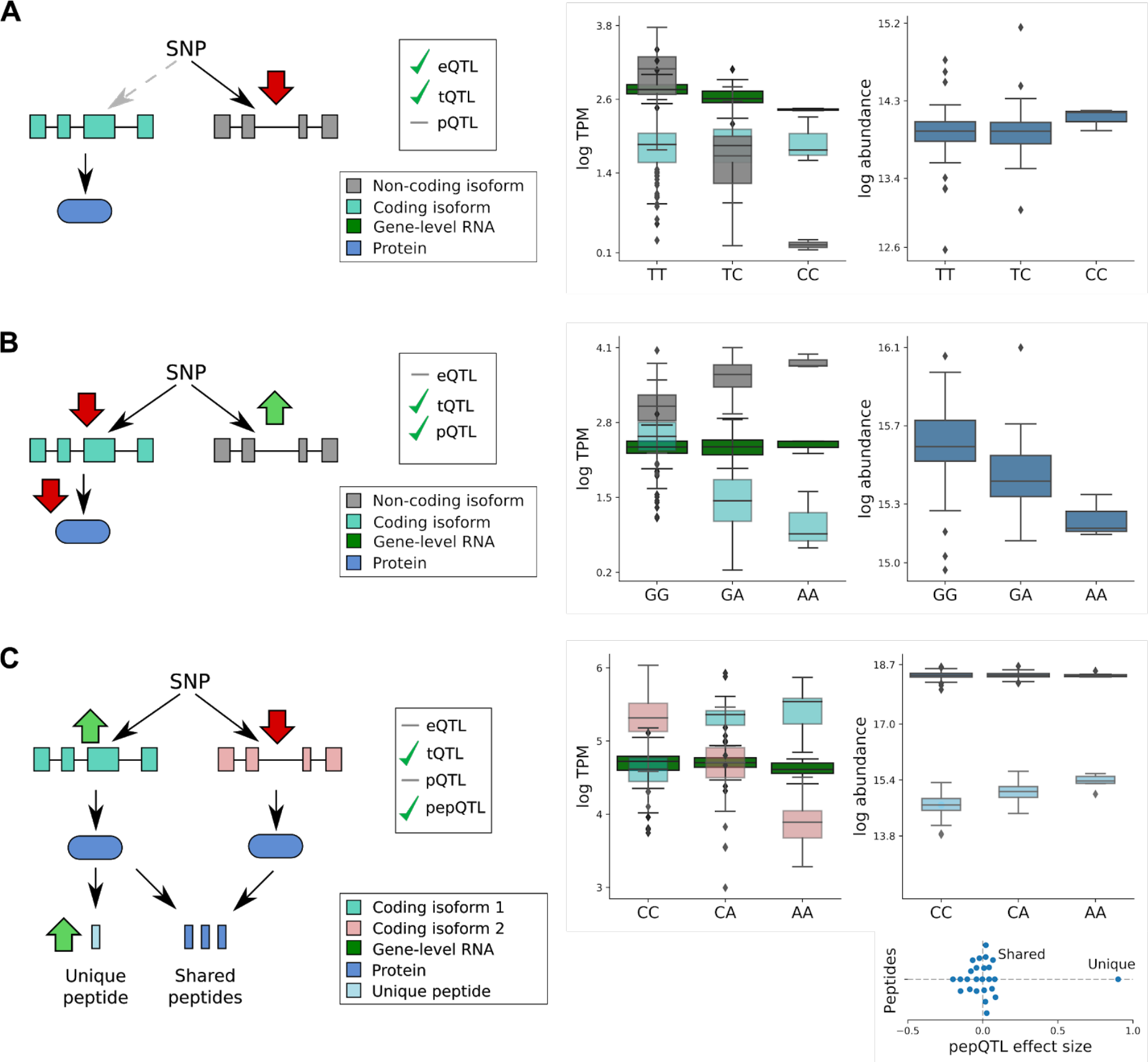
Isoform-resolution analysis of RNA and protein QTLs. **(A)** eQTL with no detectable protein effect (rs2709373; gene *METTL21A*), which can be explained by an underlying transcript QTL acting on the non-coding isoform ENST00000477919 (grey). No genetic effect is observed on the protein-coding isoform ENST00000425132 (light blue), and consequently no protein effect. **(B)** pQTL without RNA replication (rs6606721; gene *MMAB*), with a directional opposite effect on a coding and a noncoding isoform (light blue: ENST00000540016; grey: ENST00000537496), resulting in no overall change in gene expression level. **(C)** Transcript QTL that is neither an eQTL nor a pQTL. The variant rs12795503 has opposite directional effects on the two coding transcripts ENST00000301843 (light blue) and ENST00000346329 (light red), resulting in no detectable effects on either the RNA or protein level. The transcript-specific effect on ENST00000301843 is detectable for the peptide QDSAAVGFDYK (uniquely mapping to exon 11 of ENST00000301843), while no effect is observed for peptides shared by both protein isoforms. **Subplot** shows genetic effect sizes for all peptides mapped to CTTN. Shared: peptides mapping to isoforms 1 and 2; Unique: peptide uniquely mapping to isoform 1.

The transcript QTLs also provided insights into why some pQTLs were not detected as eQTLs. Out of 234 pQTLs for which no corresponding eQTL was found, we identified 66 pQTLs with a significant transcript QTL (**Supp. Fig. 13**). Interestingly, for 16 of these genes, including *MMAB* (**Fig. 3C**), we observed genetic effects with opposite directions on coding and non-coding transcript isoforms. These data show that the accurate mapping of RNA-level eQTLs can be confounded for loci that give rise to multiple transcript isoforms. In particular, transcript isoforms from the same gene may be differently affected by the same DNA variant, while only a subset of the transcripts may contribute to protein expression from the locus.

Finally, we used the peptide-level QTL information to explore, at higher resolution, isoform-specific transcript QTLs. We identified 53 genes with transcript QTLs that were not detected at either gene resolution RNA or protein levels (i.e. no eQTL or pQTL), but which were detectable as a peptide QTL. One example is the gene *CTTN* (**Fig. 4C**), where an increase in the expression of one transcript isoform was accompanied by a decrease in the expression of a second isoform. At the protein level, the same variant exerted a detectable effect on a peptide sequence that uniquely maps to the first transcript isoform.

Taken together, our results illustrate a variety of different RNA-protein relationships, and how they are affected by genetic variation between donors. These results show important roles of transcriptional regulation underlying *cis* pQTL effects, highlight mechanisms explaining the differences in observed genetic effects, and in particular show that isoform-specific effects, invisible to standard eQTL mapping approaches, can be detected at the protein level.

### *trans* protein effects of *cis* QTLs

We extended our analysis to map proteome-wide associations, considering variants with either significant *cis* RNA, or *cis* protein, QTLs (**Fig. 5A**). Overall, our data show that *cis* pQTLs have *trans* effects on protein levels more frequently than eQTLs without a corresponding *cis* pQTL (**Fig. 5B**, see **Methods**). Genome-wide we identified 89 *cis*-pQTL lead variants with *trans* effects on 173 genes (FDR<10%; **Supp. Table 15**).

**Figure 5.**
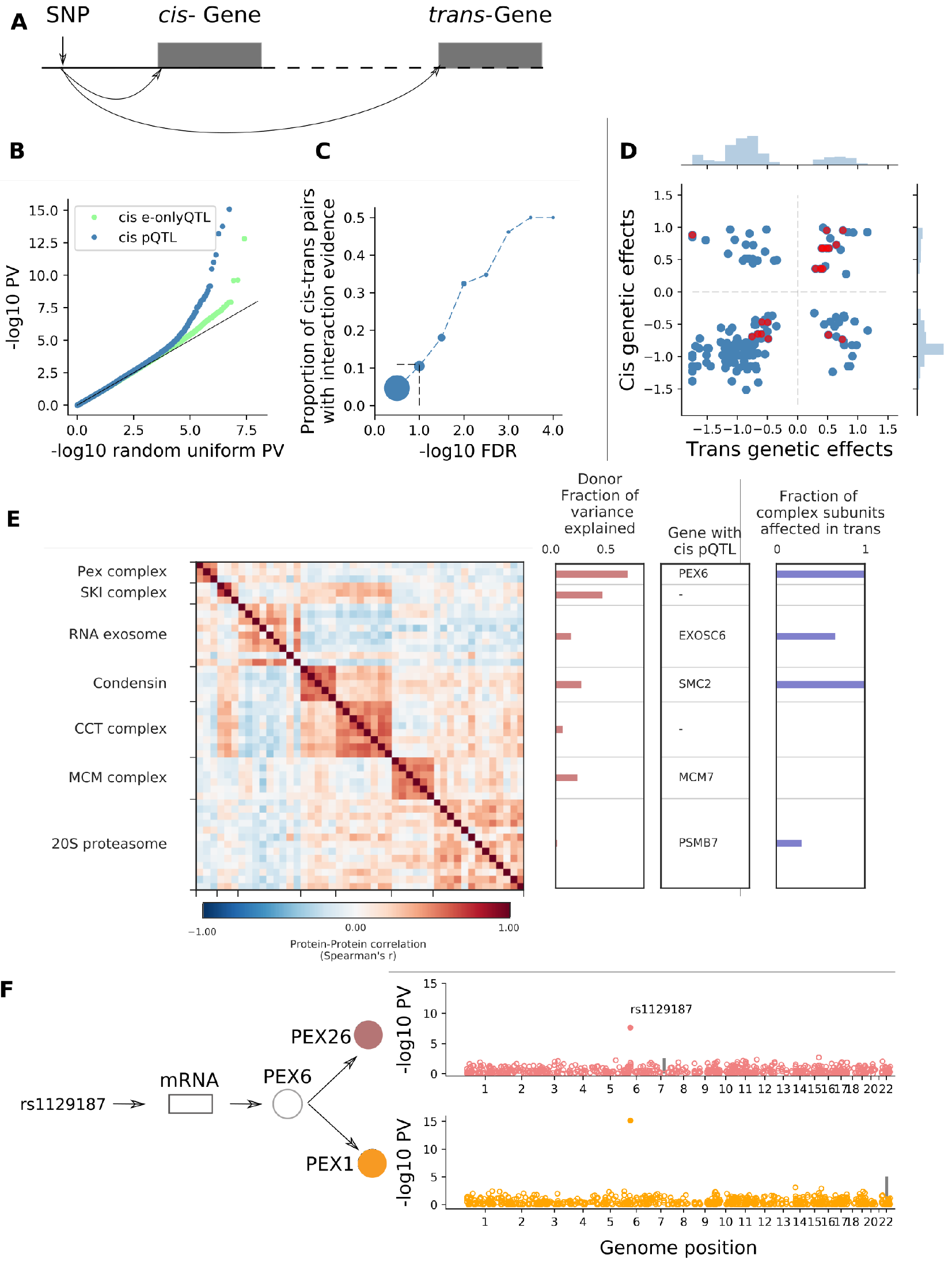
*trans* effects on the iPS proteome. **(A)** Targeted strategy for mapping *trans* genetic effect on protein abundance. Lead *cis* eQTL or pQTL variants are considered for proteome-wide association analysis. **(B)** QQ-plot of negative log P values from *trans* pQTL analysis, either considering 712 lead *cis* pQTL variants (blue) or 2,744 lead eQTL variants without replicated pQTL effect (defined as in Fig. 3A; light green) for proteome-wide association analysis. **(C)** Enrichment of protein-protein interactions among significant *trans* pQTLs. Shown is the fraction of *cis-trans* gene pairs linked by a *trans* pQTL with evidence of protein-protein interactions (based on the union of CORUM, IntAct, and StringDB), for different *trans* pQTL discovery FDR thresholds. Dot size is proportional to the number of protein pairs. Vertical line corresponds to *trans* pQTL FDR<10%. **(D)** Juxtaposition of genetic effect sizes for protein pairs that are regulated in *cis* and *trans* by the same variant (FDR<0.1). Red points indicate protein pairs with evidence for protein-protein interactions as defined in **C**. (**E**) Left: Protein coexpression of selected protein complex subunits defined based on CORUM, displaying pairwise Spearman correlation coefficients between proteins. Right: i) fraction of the average cluster protein expression level explained by donor effects; ii) subunit with the most significant *cis* pQTL; iii) fraction of subunits in association with the *cis* pQTL at nominal significance (P<0.01). (**F**) The PEX26-PEX6-PEX1 complex. The variant rs1129187 is associated in *cis* with changes in the RNA and protein abundance of PEX6 and in trans with changes in the protein abundance of PEX1 and PEX26.

We observed that groups of proteins detected with ‘shared genetic regulation’, defined here as proteins whose abundance is affected, either in *cis*, or *trans*, by the same genetic variant, were enriched for protein complex subunits (odds ratio=15, P= 1.24⋅10^−14^, Fisher’s exact test; **Fig. 5C**). The *cis* and *trans* effects showed similar effect directions and effect sizes, consistent with genetic effects mediated via stabilising protein-protein interactions (**Fig. 5D**). This hypothesis is supported by previous studies showing that protein modules sharing genetic effects in *trans* are enriched in protein interactions^23^, that somatic aberrations in human cancer cell lines are propagated in *trans*^10,11^, and by the enhanced co-expression of protein complex subunits and the significant donor variance component observed for many protein complexes (**Fig. 5E**; **Supp. Fig. 9**).

For several protein complexes, we observed that *cis* genetic regulation of one subunit may lead to *trans* genetic regulation of other subunits (**Supp. Table 16**). This is illustrated by PEX26-PEX6-PEX1, a protein complex involved in peroxisome biogenesis **(Fig. 5F).** A strong association was detected between all complex subunits and the PEX6 *cis* eQTL rs1129187 (**Fig. 3B**). This suggests that PEX6 acts as a limiting subunit of this complex in iPSCs. As noted above, this SNP is a known risk variant for Alzheimer’s disease in APOE e4+ carriers^21^. Thus, our results suggest a biological mechanism underlying this risk variant, namely through changes in the abundance of the PEX26-PEX6-PEX1 complex. This is in line with the proposed roles of peroxisomal function in the development of Alzheimer’s disease^24^.

Several variants also showed genetic effects of opposite directions in *cis* and *trans*. For example, rs1326138, the *cis* pQTL for SUCLA2, had opposite effects in *trans* on SUCLG2. These proteins are mutually exclusive binding partners of SUCLG1, with which they form the succinate coenzyme A ligase complex. A possible mechanism for this genetic effect is that an increase in SUCLA2 reduces the availability of SUCLG1 to dimerise with SUCLG2, leaving the latter in a monomeric state where it is prone to protein degradation (**Supp. Fig. 14**).

### QTLs with protein level protein level effects are enriched for human disease variants

To explore the functional physiological relevance of iPSC pQTLs, we tested for overlap with disease-linked variants identified in genome-wide association studies (GWAS). To do this, we queried QTLs that tag known GWAS variants ^25^ (i.e. are in LD r^2^>0.8; **Methods**), identifying 10% of pQTLs and 7% of eQTLs, respectively, that tag a GWAS disease variant. This corresponds to an enrichment of 1.93-fold for pQTLs and 1.36-fold for eQTLs, over a matched set of random control QTL variants (**Fig. 6A**). The data show that QTLs affecting both RNA and protein expression levels are more likely to tag a disease variant, compared with either eQTL corresponding to non-protein coding genes, or eQTL that do not result in a detectable protein effect (**Fig. 6B**). Notably, these differences could not be explained by differences in the number of eQTL and pQTL discoveries (**Supp. Fig. 15**).

**Figure 6.**
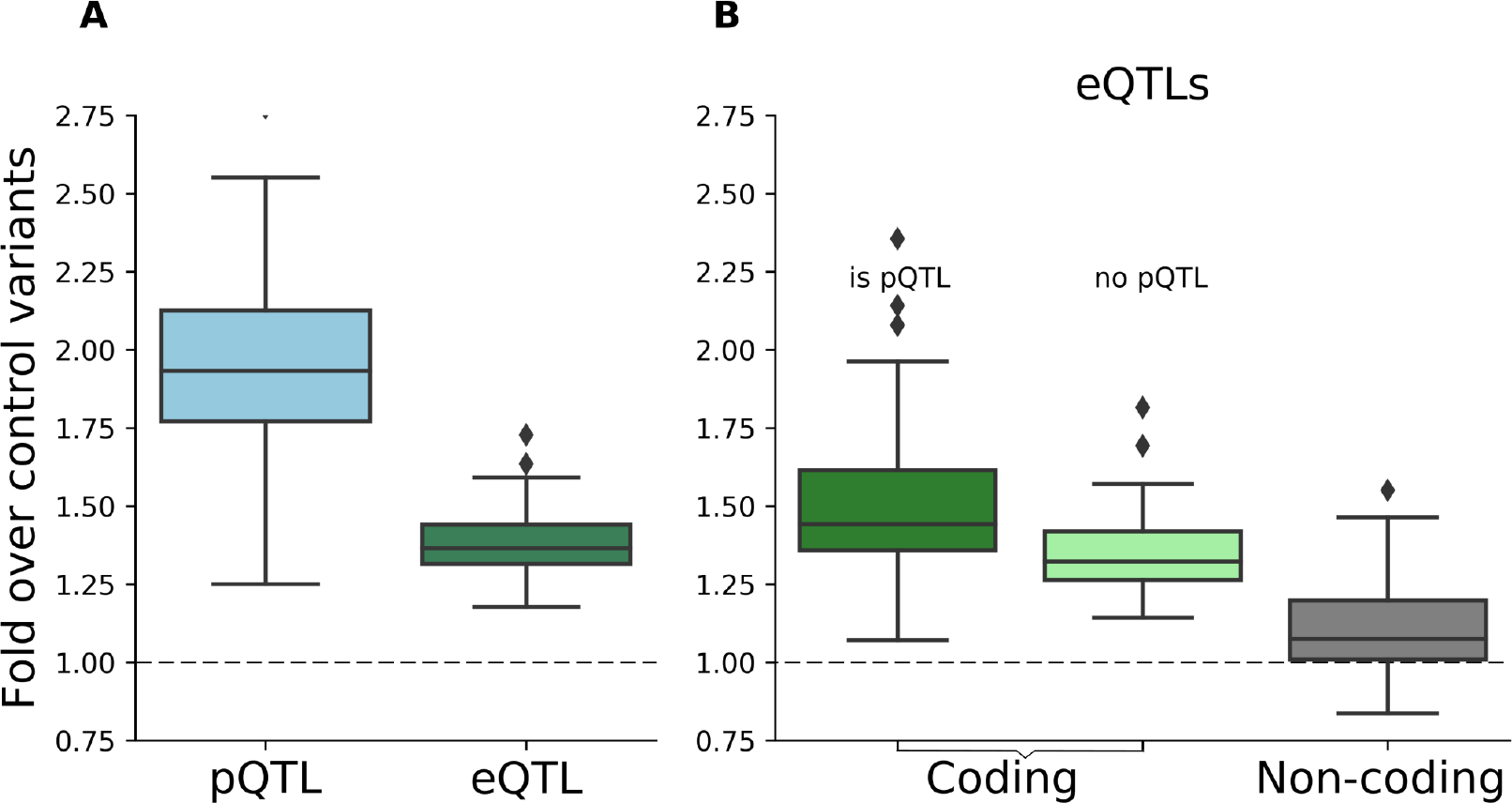
Enrichment of disease variant tagging different RNA and protein QTLs. (**A**) Fold GWAS tagging enrichment over control variants for pQTLs (blue) and eQTLs (green) corresponding to protein coding genes. (**B**) Fold enrichment for eQTLs corresponding to protein-coding genes either with (dark green) or without (light green) replicated effects at the protein level, and for eQTLs that affect non-coding genes (grey).

Of note, 19 of the pQTLs without a detectable effect at the RNA level tag GWAS variants (**Supp. Table 7**). One such example is the *cis* pQTL of VRK2, rs1051061 (**Supp. Fig. 17**), a missense variant within the kinase domain of VRK2, which is associated with schizophrenia risk^26^. VRK2, a serine/threonine kinase, is known to be down-regulated in several neurological disorders, including schizophrenia^27,28^. We hypothesise that, independently of expression changes, the alternative allele of rs1051061 affects the protein structure and its capacity to bind, leaving the protein in an unstable state. This result contributes to the understanding of schizophrenia’s aetiology, supporting an important role for VRK2 and suggesting possible disease onset already in early development stages, i.e in pluripotent cells.

In summary, our data strongly support the conclusion that analysis of pQTLs provides unique information regarding the functioning of disease risk variants and give insights, which are not identifiable using eQTL mapping, into mechanisms through which genetic effects modulate cell physiology.

## Discussion

We have performed the first in-depth characterisation of gene expression and the human iPSC proteome, and, to our knowledge, provided the largest dataset with parallel RNA/protein profiling in human cells. By quantifying protein and transcript expression variation across more than 200 human iPSC lines, we identified genetic and non-genetic mechanisms underlying variation at both the protein and RNA levels. We have mapped more than 700 protein Quantitative Trait Loci (pQTLs) and analysed in detail how these relate to eQTLs. While previous studies have established overlap and colocalization of eQTL and disease-linked GWAS associations^29^, a key finding from this study is that the subset of QTLs with an effect at the protein level were significantly more likely to be associated to disease traits. These results demonstrate the importance of the systematic identification of mechanisms through which genetic variation can affect cell physiology and disease.

We have identified the specific proteins that show most variation in abundance between iPSC lines. These are often co-expressed in groups of proteins with shared biological functions. Thus, the major variation is seen with proteins affecting processes such as cell differentiation and cell-cell adhesion. Importantly, we detected many proteins that varied in abundance without a parallel variation in the abundance of their cognate RNAs. These observations indicate an important role for post-transcriptional mechanisms in contributing to genetic variation in the human population and identify genes whose important roles are invisible in transcript mapping studies.

Our data identified that donor-specific genetic factors were major contributors to the differences in protein expression detected across the iPSC lines. Another major factor was the cell culture conditions, indicating that protein expression in iPSCs is sensitive to the cellular growth environment. Consistent with the significant influence of donor genetics on variation in protein expression, we mapped 712 common genetic variants associated with changes in protein abundance. By the systematic comparison of matched protein and RNA data, including detailed resolution of separate isoforms, we identified that in *cis*, DNA variants act mainly through transcriptional mechanisms. This involves the variant either modulating total transcript abundance, or, in some cases, varying the proportions of different transcript isoforms produced from the locus. This extends previous results on the strong overlap between *cis* eQTLs and pQTLs^7^.

Our data also illustrate the ability of protein-protein interactions to both buffer and propagate genetic effects. A long-standing hypothesis has been that many protein complexes have a rate-limiting subunit that determines complex abundance, with any excess subunits being rapidly degraded (e.g. because of exposure of hydrophobic residues). This has two implications. First, *cis* eQTLs for non-rate-limiting subunits should have minimal effect at the protein level, since the abundance of these proteins is determined by the abundance of the whole complex. Second, *cis* eQTLs for rate-limiting subunits should have effects in *trans* on the abundance of the whole complex, and on most, if not all, subunits therein. We found evidence for both phenomena in our analysis of *cis* and *trans* pQTLs. These observations, the first to our knowledge for common genetic variants in human, are consistent with previous results obtained on high heterozygosity samples, i.e outbred mice^23^, and somatic aberrations in human cancer cell lines^10,11^.

Understanding the mechanisms through which genetic variations act in the human population is of great relevance to characterising risk factors and susceptibility to disease. There is growing interest in the potential for studying disease mechanisms using disease relevant tissues that are derived from panels of iPSCs^30–33^. Our study provides important information for advancing such studies on the genetic regulation of protein expression and disease-relevant phenotypes in iPSC-derived model systems.

## Author contributions

DS, BM, AL, OS: Wrote the paper with input from all authors.

DS, BM, AB, HK, MB: Contributed to the bioinformatic analysis

DB: Generated the proteomics data

MB: Curated and processed the RNA data

AB, BM, DB: Curated and processed proteomics data

DS: Analysed the data - variance component analysis

BM: Analysed the data - QTL analysis

DS, MB, BM: Designed the QTL mapping workflow

AL, OS: Supervised and designed the research.

## Acknowledgements

This work was funded with a strategic award from the Wellcome Trust and Medical Research Council (WT098503). The authors thank our colleagues, including Pedro Beltrao, Doreen Cantrell, Jason Swedlow and Greg Findlay, for comments on the manuscript.

## Methods

### RNA-seq data processing

Raw RNA-seq data for 331 samples were obtained from the ENA project: ERP007111. CRAM files were merged on a sample level and were converted to FASTQ format. The reads were trimmed to remove adapters and low quality bases (using Trim Galore!^34^), followed by read alignment using STAR (version: 020201)^35^, using the two-pass alignment mode and the default parameters as proposed by ENCODE (c.f. STAR manual). All alignments were relative to the GRCh37 reference genome, using ENSEMBL 75 as transcript annotation^36^.

Samples with low quality RNA-seq were discarded, if they had less than 2 billion bases aligned, had less than 30% coding bases, or had a duplication rate higher than 75% were. This resulted in 323 lines for analysis, for 202 of which matched proteome data was available.

Gene-level RNA expression was quantified from the STAR alignments using featureCounts (v1.6.0)^37^, which was applied to the primary alignments using the “-B” and “-C” options in stranded mode, using the ENSEMBL 75 GTF file. Quantifications per sample were merged into an expression table using the following normalization steps. First, gene counts were normalized by gene length. Second, the counts for each sample were normalized by sequencing depth using the edgeR adjustment^38^.

Transcript isoform expression was quantified directly from the (unaligned) trimmed reads using Salmon^22^ (version: 0.8.2), using the “--seqBias”, “--gcBias” and “VBOpt” options in “ISR” mode to match our inward stranded sequencing reads. The transcript database was built on transcripts derived from ENSEMBL 75. The TPM values as returned by Salmon were combined into an expression table

### Quantitative proteomics data generation

All lines included in this study are part of the HipSci resource and were reprogrammed from primary fibroblasts as previously described^1^. We selected 217 lines for in depth proteomic analysis with Tandem Mass Tag Mass Spectrometry. A subset of 202 lines (112 normal and 90 disease; **Supp. Table 1**) with matched mRNA and protein data were considered for further analysis.

#### Sample preparation

For protein extraction, frozen iPSC cell pellets were washed with ice cold PBS and redissolved immediately in 200 μL of lysis buffer (8 M urea in 100 mM triethyl ammonium bicarbonate (TEAB) and mixed at room temperature for 15 minutes. DNA in the cell lysates was sheared using ultrasonication (6 × 20 s at 10°C). The proteins were reduced using tris-carboxyethylphosphine TCEP (25 mM) for 30 minutes at room temperature, then alkylated in the dark for 30 minutes using iodoacetamide (50 mM). Total protein was quantified using the fluorescence based EZQ assay (Life Technologies). The lysates were diluted 4-fold with 100 mM TEAB for the first protease digestion with mass spectrometry grade lysyl endopeptidase, Lys-C (Wako, Japan), then diluted a further 2.5-fold before a second digestion with trypsin. Lys-C and trypsin were used at an enzyme to substrate ratio of 1:50 (w/w). The digestions were carried out for 12 hours at 37°C, then stopped by acidification with trifluoroacetic acid (TFA) to a final concentration of 1% (v:v). Peptides were desalted using C18 Sep-Pak cartridges (Waters) following manufacturer’s instructions and dried.

#### Tandem Mass Tag Mass Spectrometry analysis

For Tandem Mass Tag (TMT)-based quantification, the dried peptides were redissolved in 100mM TEAB (50 μL) and their concentration was measured using a fluorescent assay (CBQCA) (Life Technologies). 100 μg of peptides, from each cell line to be compared, in 100 μL of TEAB were labelled with a different TMT tag (20 μg ml^−1^ in 40 μL acetonitrile) (Thermo Scientific), for two hours at room temperature. After incubation, the labelling reaction was quenched using 8 μl of 5% hydroxylamine (Pierce) for 30 minutes and the different cell lines/tags were mixed and dried in vacuo. TMT-ten plex was used to label ten iPSC lines and quantify them in parallel. In total 24 TMT-ten plex experiments were performed, where one iPSC line (bubh_3) was chosen as a reference cell line and was kept constant in all TMT batches. The other nine quantification channels were used to label 9 different cell lines.

The TMT samples were fractionated using off-line high pH reverse phase chromatography: samples were loaded onto a 4.6 × 250 mm Xbridge™ BEH130 C18 column with 3.5 μm particles (Waters). Using a Dionex bioRS system, the samples were separated using a 25-minute multistep gradient of solvents A (10 mM formate at pH 9) and B (10 mM ammonium formate pH 9 in 80% acetonitrile), at a flow rate of 1 ml/min. Peptides were separated into 48 fractions, which were consolidated into 24 fractions. The fractions were subsequently dried and the peptides redissolved in 5% formic acid and analysed by LC-MS.

5% of the material was analysed using an orbitrap fusion tribrid mass spectrometer (Thermo Scientific), equipped with a Dionex ultra high-pressure liquid chromatography system (nano RSLC). RP-LC was performed using a Dionex RSLC nano HPLC (Thermo Scientific). Peptides were injected onto a 75 μm × 2 cm PepMap-C18 pre-column and resolved on a 75 μm × 50 cm RP-C18 EASY-Spray temperature controlled integrated column-emitter (Thermo), using a four-hour multistep gradient from 5% B to 35% B with a constant flow of 200 nL min^−1^. The mobile phases were: 2% ACN incorporating 0.1% FA (Solvent A) and 80% ACN incorporating 0.1% FA (Solvent B). The spray was initiated by applying 2.5 kV to the EASY-Spray emitter and the data were acquired under the control of Xcalibur software in a data dependent mode using top speed and 4 s duration per cycle. The survey scan is acquired in the orbitrap covering the *m/z* range from 400 to 1400 Th, with a mass resolution of 120,000 and an automatic gain control (AGC) target of 2.0 e5 ions. The most intense ions were selected for fragmentation using CID in the ion trap with 30 % CID collision energy and an isolation window of 1.6 Th. The AGC target was set to 1.0 e4 with a maximum injection time of 70 ms and a dynamic exclusion of 80 s.

During the MS3 analysis for more accurate TMT quantifications, 5 fragment ions were co-isolated using synchronous precursor selection using a window of 2 Th and further fragmented using HCD collision energy of 55% ^39^ The fragments were then analysed in the orbitrap with a resolution of 60,000. The AGC target was set to 1.0 e5 and the maximum injection time was set to 105 ms.

### Proteomics data processing

The TMT labeled samples (24 batches of TMT-ten plex) were analysed using MaxQuant v. 1.6.0.13 ^40,41^. Proteins and peptides were identified using the UniProt *human* reference proteome database (Swiss Prot + TrEMBL) release-2017_03, using the Andromeda search engine. Run parameters and the raw MaxQuant output have been deposited at PRIDE (PXD010557).

The following search parameters were used: reporter ion quantification, mass deviation of 6 ppm on the precursor and 0.5 Da on the fragment ions; Tryp/P for enzyme specificity; up to two missed cleavages, “match between runs”, “iBAQ”. Carbamidomethylation on cysteine was set as a fixed modification. Oxidation on methionine; pyro-glu conversion of N-terminal Gln, deamidation of asparagine and glutamine and acetylation at the protein N-terminus were set as variable modifications^40–42^.

Peptides and protein groups were identified at a False Discovery Rate (FDR) of 5%. The same FDR was applied to the Post-Translational Modifications (PTM) Site and the Peptide Spectrum Matches (PSM). We performed the FDR calculation on an extended set and removed the Razor Protein FDR calculation constrain (for more details see reference ^43^). In total we identified 255,015 peptides detected in at least one sample (after removing reverse and contaminant peptides; on the 217 lines and 23 replicates of the reference line), which corresponds to 16,773 protein groups.

#### Quality control and quantification

To rule out technical confounding when performing genetic analyses of protein traits, we discarded 2,072 peptides that overlap a non-synonymous common variant (MAF>5% in European population) in expressed transcript (average TPM>1 based on RNA-seq). Protein group abundances were then estimated as the sum of peptide intensities mapped to a protein group. For peptide abundance we use the intensities reported in the “Peptides” file from MaxQuant.

We discarded 10 lines with fewer than 67,000 identified peptides (corresponding to %75 of the median number of peptides identified; **Supp. Fig. 1**), resulting in a proteomics dataset consisting of 207 lines, 202 of which had matched RNA-Seq data and hence were considered for further analysis. In addition, the technical replicates for the included reference line in each TMT batch were retained to aide the normalization of protein quantifications between batches; see below.

In aggregate across all lines, we detected 16,218 protein groups. For downstream analysis, we considered protein groups that were detected in at least 30 of the 202 lines and analogously considered recurrently detected peptides (**Supp. Fig. 2**), resulting in a final dataset of 11,542 recurrent protein groups and 132,716 recurrent peptides. These protein groups could be mapped to 9,993 protein coding genes.

To adjust for technical effects during the acquisition of protein data in TMT batches, we scaled the abundance estimate for each feature (i.e protein or peptide) as follows. For a feature and TMT batches, a scaling coefficient was computed as the ratio between the median intensity value across all lines versus the median intensity value across the subset of lines within the batch.

Next, we employed quantile normalization across the feature abundance distribution in each line, using a normalization reference line (selected as the line with the highest number of total peptides detected), Briefly, for each line and feature we replaced the observed expression value with the expression level in the reference line having the same rank position in the line to be normalized: 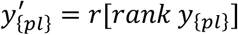, where y_{pl}_ are the intensity values for feature p and line l obtained after batch scaling, i.e. before normalisation, *r* is the sorted vector of intensities from the normalisation reference line, and 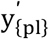 is the normalized value.

Following the approach in ^7^, we assessed quantitative compression in our proteomics data by examining changes in peptides overlapping non-synonymous variants. A non-synonymous variant in a peptide prevents detection of that peptide, as its sequence will not exist in the proteome reference. Thus, in samples heterozygous for the non-synonymous variants, the measured peptide abundance is expected to be half of that of samples homozygous for the reference variant. Our data are consistent with this expectation, indicating that compression effects are minimal in our study (**Supp Fig. 12**).

### Comparisons of iPS proteome profiles to existing tissue datasets

In order to compare our iPSC proteome dataset to the Human Proteome Map (HPM) ^15^ (**Fig. 1D**), we first mapped the RefSeq IDs of proteins quantified in the HPM to UniProt IDs. We then considered the subset of 8,333 proteins with mappable IDs that were expressed in our iPSC dataset and in at least one HPM tissue. We then calculated spearman correlation coefficients between the aggregate iPS proteome abundance profile (averaged across lines) and each HPM tissue.

### RNA-protein correlations

For global correlations of RNA and protein abundance across all genes (**Fig 1C**), the mean abundance of each RNA and protein (using TPM and iBAQ scales, respectively) was calculated across all samples, and then the Spearman correlation across all RNA-protein pairs. For correlations of RNA and protein abundance across samples for each gene (**Fig 2B**), Pearson’s correlations were calculated on the subset of samples for which both RNA and protein data were available (i.e. there no imputation or substitution of zeros for missing values in the protein data). In both cases, multi-mapping IDs between RNAs and proteins were resolved by choosing one mapping at random, dropping multi-mapping IDs from the set of protein IDs first, then from the set of gene IDs.

### RNA and protein variance component analysis

In order to calculate the contribution of each factor k to variation in protein abundance, we fitted a random effects model: 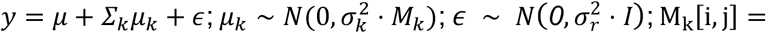 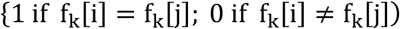. Here y denotes the (N x 1) vector of log-scaled protein abundances (or, for a coexpression cluster, the log-scaled median abundance of proteins in the cluster), *μ*_*k*_ are the random effects, *M*_*k*_ is the (N x N) covariance structure, *σ*_*k*_ is the standard deviation, and *ε* is the residual (i.i.d. noise). The random effect components are defined based on a categorical covariance function defined on covariates *f*_*k*_, that is the vector of observed values for factor k (e.g. *f*_*k*_[*i*] ∈ {*′male′, ′female′*} when k is the donor sex component). We considered donor identity, donor sex, donor age, culture medium, TMT batch, and TMT channel as random effect components. In order to accurately estimate donor variance component, we restricted this analysis to the set of lines from the subset of 51 donors for which 2 cell lines were assayed. Analogous analyses were considered for RNA abundance, leaving out the TMT-specific random effects.

In order to account for the effects of RNA abundance on protein abundance, we also applied the variance decomposition analysis to protein abundance values after adjusting for RNA variation. Adjusted protein abundances were calculated by regressing out the effects of RNA abundance (i.e. gene-level quantifications of RNA) on protein abundance for each RNA-protein pair. To do this, we fitted a linear model between RNA and protein abundances across lines (using the Numpy function poly1d in Python), taking the model residuals as the adjusted protein abundance values. Variance decomposition models were then fitted as described above.

All variance component models were fitted using the LIMIX package^44^ (https://github.com/limix/limix).

### Protein co-expression and GO enrichment analysis

Proteins were clustered into groups based on their patterns of coexpression. Coexpression was quantified by the Spearman correlation (r) between pairs of proteins. Clustering was performed using the affinity propagation algorithm ^45^, as implemented in the scikit-learn python library, with the preference parameter (determining the number of clusters identified) set to −5.0 for protein, and the damping parameter set to 0.8. Median expression of each cluster in each line was calculated by mean-normalising each protein (i.e. setting mean abundance across all samples for each protein to 1), and taking the median across all proteins in each cluster in each sample. GO enrichments for each cluster were computed using the goatools package (https://github.com/tanghaibao/goatools), and are provided in **Supp. Table 5**.

### QTL mapping of RNA and protein traits

#### *cis* QTL mapping

We used PEER ^20^ to account for unwanted variation and confounding factors both for RNA and protein traits. PEER was applied to log normalized protein abundance and log normalized gene TPM, considering the most highly expressed 10,000 proteins and genes, respectively. We selected 7 factors for protein and 13 factors for RNA, settings that were determined as the largest number of uncorrelated PEER factors identified (r<0.7; **Supp. Fig. 10**).

At protein level (protein and peptide traits), we considered the subset of lines with non-zero abundance for analysis. For RNA (gene and transcript isoform traits) all analyses are based on data from all 202 lines.

For *cis* genetic analyses, we considered common variants (MAF>5%) in gene-proximal regions of 250k upstream and downstream of gene transcription start and end sites (GRCh37). We used a linear mixed model implemented in LIMIX ^44^, to control for both population structure and repeat lines from the same donor using kinship as a random effect component. The population structure random effect component was estimated as the realized relationship covariance, i.e. dot product of the genotype matrices. PEER factors were included as fixed effect covariates in all analyses.

We used an approximate permutation scheme as in Fast QTL ^46^, based on a parametric fit to the null distribution, to adjust for multiple testing across *cis* variants for each gene. Briefly, for each gene, we obtained p-values from 100 permutations of *cis* variants. We then estimated an empirical null distribution by fitting a parametric Beta distribution to the obtained p-values. Using this null model, we estimated *cis* region adjusted p-values for QTL lead variants. For multiple testing adjustment across genes, we performed Benjamini-Hochberg adjustment. This procedure was applied to perform *cis* eQTL mapping.

For protein, peptide and transcript QTLs, herein features, we reported results at gene level and accounted for multiple testing across features mapping to the same gene. Subsequent to the permutation-based adjustment for individual features per gene, we applied a Bonferroni correction to the *cis* region adjusted p-values. We then identify the lead QTL variant and feature at the gene level, i.e. the combination of the most associated variant and trait (*cis* region and across features adjusted). The Benjamini-Hochberg procedure was applied on the gene level lead QTLs for adjustment across genes.

#### *trans* QTL mapping

*Trans* QTLs mapping was applied in a targeted manner, considering lead cis QTLs (712 pQTLs and 2,744 eQTLs not replicated at pQTL level; FDR <10%), testing each of the 11,542 recurrently expressed proteins. Genome-wide Benjamini Hochberg adjustment was performed across all tests (8⋅10^6^ variants × proteins for *cis* pQTLs).

### Downstream analysis of QTL results

#### QTL replication

We defined a lead QTL variant as ‘replicating’ across molecular layers if it had, for the same gene, a statistically significant effect and the same direction of effect on both layers. For the replicating layer, the statistical significance is defined using the nominal p-value (P<0.01), or the Bonferroni corrected value (P<0.01/N, where N is the number of features) if multiple features map to the same gene.

#### *cis* eQTL and pQTL replication

We trained a multivariate logistic regression model to the replication status of 1,887 genes with an eQTL for which the protein and the RNA were identified in all lines (**Supp. Table 13**). This stringent filter on the set of genes was used to mitigate effects due to differences in samples size (pQTLs tests were performed on the set of in which the protein was detected). For each RNA-protein pair, we defined 7 factors. The “protein coefficient of error” factor was computed as the coefficient of variation across the set of technical replicates (i.e. across the replicate measurements of the reference sample that was included in every TMT batch). The “protein complex membership” factor was assessed using existing annotation (CORUM release May 2017; ^47^), which was set to one if the gene encodes for the subunit of a protein complex and zero otherwise. The “only-nc-tQTLs” factor was obtained by assessing the replication of eQTLs for protein coding genes in transcript isoform QTLs (tQTLs), which was set to one if the eQTL was replicated in tQTL corresponding to a non-protein coding transcript isoform coding tQTL (but not in one corresponding to a protein coding isoform). When this assessment was not possible, or when the eQTL was replicated in at least one coding tQTL, we set the factor to 0.

We enabled comparison across factors by binarizing the values for eQTL effect size, average protein abundance, protein coefficient of variation across lines, the number of peptide identified for each protein, and protein coefficient of error. The factor was considered to be present for values higher than the mean across all genes and zero otherwise.

#### Annotation of *cis-trans* protein pairs with protein-protein interactions

Protein-protein interactions were obtained from the union of CORUM ^47^, IntAct ^48^ and protein-binding interactions from StringDB ^49^. In CORUM, we considered pairwise interactions between all protein complex subunits. When assessing the consistency of cis-trans pQTL paris, we discarded any isoform extension from the protein UniProt IDs and intersected the gene pair with the aggregate protein-protein interactions reference list.

#### Overlap with disease variants

Following the approach in ^1^, we defined proxy variants of each cis QTL as variants in high LD (r^2^ > 0.8; based on the UK10K European reference panel50) within the same cis window. A QTL was defined as GWAS-tagging if at least one such proxy variant was annotated in the NHGRI-EBI Gwas catalog (download on 10 April 2018; converted to hg19). We considered a stringent subset of 21,601 associations for analysis (out of 65,761 total associations), that were i) genome-wide significant (P<5⋅10-8) and ii) reported in studies with a sample sizes of at least 1,000, individuals, and iii) for which the effect size (odds ratio) was reported in the catalogue.

To assess the enrichment of different QTL types for GWAS variants, we compared the fraction of GWAS-tagging QTL variants to sets of random matched control variants that were drawn from the European 1000G phase 3 ^50^, matched for minor allele frequency, the number of variants in LD (‘LD buddies’; r^2^ > 0.5), distance to the nearest gene, and gene density, allowing for maximum deviation of +/− 50% for each criterion. For each QTL type, we generated 100 sets of control variants using SNPsnap^51^, based on the respective QTL variants as the input.

## Data availability

RNA-Seq data for 331 samples are available on the European Nucleotide Archive (ENA): study PRJEB7388; accession ERP007111. Proteomics quantifications (protein group and peptide resolution; MaxQuant output), and run parameters will be available on the PRIDE Archive (PXD010557).

